# Evolutionary stability of collateral sensitivity to antibiotics in the model pathogen *Pseudomonas aeruginosa*

**DOI:** 10.1101/570663

**Authors:** Camilo Barbosa, Roderich Roemhild, Philip Rosenstiel, Hinrich Schulenburg

## Abstract

Evolution is at the core of the impending antibiotic crisis. Sustainable therapy must thus account for the adaptive potential of pathogens. One option is to exploit evolutionary trade-offs, like collateral sensitivity, where evolved resistance to one antibiotic causes hypersensitivity to another one. To date, the evolutionary stability and thus clinical utility of this trade-off is unclear. We performed a critical experimental test on this key requirement, using evolution experiments with *Pseudomonas aeruginosa* combined with genomic and genetic analyses, and identified three main outcomes: (i) bacteria commonly failed to counter hypersensitivity and went extinct; (ii) hypersensitivity sometimes converted into multidrug resistance; and (iii) resistance gains occasionally caused re-sensitization to the previous drug, thereby maintaining the trade-off. Drug order affected the evolutionary outcome, most likely due to variation in fitness costs and epistasis among adaptive mutations. Our finding of robust genetic trade-offs and drug-order effects can guide design of evolution-informed antibiotic therapy.

## Introduction

Treatment of infectious diseases and cancer often fail because of the rapid evolution of drug resistance (Bloemberg et al., 2015; Davies & Davies, 2010; Gottesman, 2002; Zaretsky et al., 2016). Optimal therapy should thus anticipate how resistance to treatment evolves and exploit this knowledge to improve therapy (Gatenby, Silva, Gillies, & Frieden, 2009; Imamovic & Sommer, 2013). One promising strategy is based on evolved collateral sensitivity: the evolution of resistance against one drug A concomitantly causes hypersensitivity (i.e., collateral sensitivity) to a second drug B (Szybalski & Bryson, 1952). If evolved collateral sensitivity is reciprocal, it can – in theory – trap the bacteria in a double bind, thereby preventing the emergence of multi-drug resistance during treatment (Baym, Stone, & Kishony, 2016; Pál, Papp, & Lázár, 2015; Roemhild & Schulenburg, 2019). Recent large-scale studies have demonstrated that evolved collateral sensitivity is pervasive in laboratory strains and clinical isolates of distinct bacterial species (Barbosa et al., 2017a; Imamovic et al., 2018; Imamovic & Sommer, 2013; Jansen et al., 2016; Jiao, Baym, Veres, & Kishony, 2016; Lázár et al., 2014, 2013; Oz et al., 2014; Podnecky et al., 2018) as well as cancer cells (Dhawan et al., 2017; Pluchino, Hall, Goldsborough, Callaghan, & Gottesman, 2012; Shaw et al., 2015; Zhao et al., 2016). More importantly, evolved collateral sensitivity can slow down resistance evolution during combination (Barbosa, Beardmore, Schulenburg, & Jansen, 2018; Evgrafov, Gumpert, Munck, Thomsen, & Sommer, 2015; Munck, Gumpert, Wallin, Wang, & Sommer, 2014) and sequential therapy (Kim, Lieberman, & Kishony, 2014; Roemhild, Barbosa, Beardmore, Jansen, & Schulenburg, 2015), and also limit the spread of plasmid-borne resistance genes (Rosenkilde et al., 2019).

Although collateral sensitivity appears to be pervasive, its utility for medical application is still dependent on several additional factors. Firstly, the evolution of collateral sensitivity should ideally be repeatable for a given set of conditions (Nichol et al., 2019). This means that independent populations selected with the same drug should produce identical collateral effects when exposed to a second one. Such high repeatability is not always observed. Recent work even identified evolution of contrasting collateral effects (i.e., some populations with evolved collateral sensitivity and others with cross-resistance) for different bacteria, including *Pseudomonas aeruginosa* (Barbosa et al., 2017), *Escherichia coli* (Nichol et al., 2019; Oz et al., 2014), *Enterococcus faecalis* (Maltas & Wood, 2019), and a BCR-ABL leukemeia cell line (Zhao et al., 2016). These patterns are likely due to the stochastic nature of mutations combined with alternative evolutionary paths to resistance against the first selective drug, subsequently causing distinct collateral effects against other drugs (Barbosa et al., 2017a; Nichol et al., 2019). Secondly, the evolution of collateral sensitivity should ideally be repeatable across conditions, for example different population sizes. This is not always the case. For example, an antibiotic pair, which consistently produced collateral sensitivity in small *Staphylococcus aureus* populations (e.g., 10^6^), instead produced complete cross-resistance in large populations (e.g., 10^9^) and thus an escape from the evolutionary constraint, most likely due to the higher likelihood of advantageous rare mutations under these conditions (Jiao et al., 2016).

A third and largely unexplored factor is that evolved collateral sensitivity and, hence, the resistance trade-off should be stable across time. This implies that bacteria either cannot evolve to overcome collateral sensitivity and thus die out, or, if they achieve resistance to the new drug B, they should concomitantly be re-sensitized to the original drug A. Two recent studies, both with different main research objectives, yielded some insight into this topic. One example was focused on historical contingency during antibiotic resistance evolution of *P. aeruginosa* (Yen & Papin, 2017). As part of the results, the authors identified lineages with evolved resistance against ciprofloxacin that simultaneously showed increased sensitivity to piperacillin and tobramycin. The reverse pattern (i.e., evolved high resistance to either piperacillin or tobramycin and increased sensitivity to ciprofloxacin) was not observed and, thus, this case represents an example of uni-directional collateral sensitivity. The subsequent exposure of the ciprofloxacin-resistance lineages to either piperacillin or tobramycin led to the evolution of resistance against these two antibiotics and substantial (yet not complete) re-sensitization to ciprofloxacin. The second study focused on evolving *E. coli* populations in a morbidostat, in which the bacteria were exposed to repeated switches between two drugs (Yoshida et al., 2017). The evolution of multi-drug resistance was only prevented in the two treatments with polymyxin B that were also characterized by evolved collateral sensitivity, although again only in one direction (Yoshida et al., 2017). To date, the general relevance of this third factor is still unclear, especially for conditions when collateral sensitivity is reciprocal and when the evolving populations are also allowed to go extinct (i.e., they cannot overcome the evolutionary trade-off).

Here, we specifically tested the potential of the model pathogen *P. aeruginosa* to escape reciprocal collateral sensitivity through *de novo* evolution. We focused on the first switch between two drugs, because the evolutionary dynamics after this first switch will reveal the ability of the bacteria to adapt to the second drug, against which they evolved sensitivity, and, if so, whether this causes re-sensitization to the first drug. These two aspects are key criteria for applicability of a treatment strategy that exploits evolved collateral sensitivities. Our analysis is based on a two-step evolution experiment. Bacteria first evolved resistance against a first drug A and concomitant sensitivity against a second drug B. Thereafter, bacteria were subjected to a second evolution experiment, during which they were allowed to adapt to the second drug B, either alone or additionally in the presence of the first drug A. Phenotypic characterization of the evolved bacteria was combined with genomic and functional genetic analyses, in order to determine the exact targets of selection under these conditions.

## Results

The experimental design of our two-step evolution experiment is illustrated in Fig. 1a. We took advantage of previously evolved, highly resistant *P. aeruginosa* populations, which we obtained from serial-passage experiments with increasing concentrations of clinically relevant bactericidal antibiotics (drug A, Fig. 1a) (Barbosa et al., 2017). From these, we identified two cases of reciprocal collateral sensitivity, including (i) piperacillin/tazobactam (PIT) and streptomycin (STR), or (ii) carbenicillin (CAR) and gentamicin (GEN). In the current study, we now re-assessed the reciprocity of collateral effects using dose-response analysis (Fig. 1b and c – Source Data 1). Thereafter, we isolated resistant colonies from these populations and switched treatment to the drug, against which collateral sensitivity was observed (drug B, Fig. 1a). The evolutionary challenge was initiated at sub-inhibitory concentrations of each drug (vertical black dashed lines in Fig. 1b and c), followed by linear concentration increases at two different rates: mild or strong (vertical orange and blue dashed lines respectively, Fig. 1b and c). We specifically chose linear increases, because our main objective was to better understand the evolutionary dynamics occurring during the first switch of a collateral sensitivity cycling strategy. Linear increases would, in this case, facilitate evolutionary rescue and provide ample opportunity to escape the double bind, thereby yielding a conservative measure for the applicability of collateral sensitivity as a treatment strategy. We additionally considered treatments where antibiotics were also switched to collateral sensitivity, but selection by drug A was continued in combination with drug B; hereby denoted as constrained environments. Overall, four selective conditions were run in parallel: mild or strong increases of the second drug B in either the presence (constrained) or absence (unconstrained) of the first drug A (Fig. 1a). We further included control experiments without antibiotics. To determine treatment success, we monitored bacterial growth with continuous absorbance measurements, quantified frequencies of population extinction, and characterized changes in antibiotic resistance of the evolved bacteria as previously evaluated for *P. aeruginosa* and other bacteria (Barbosa et al., 2018, 2017; Hegreness, Shoresh, Damian, Hartl, & Kishony, 2008).

**Figure 1.**
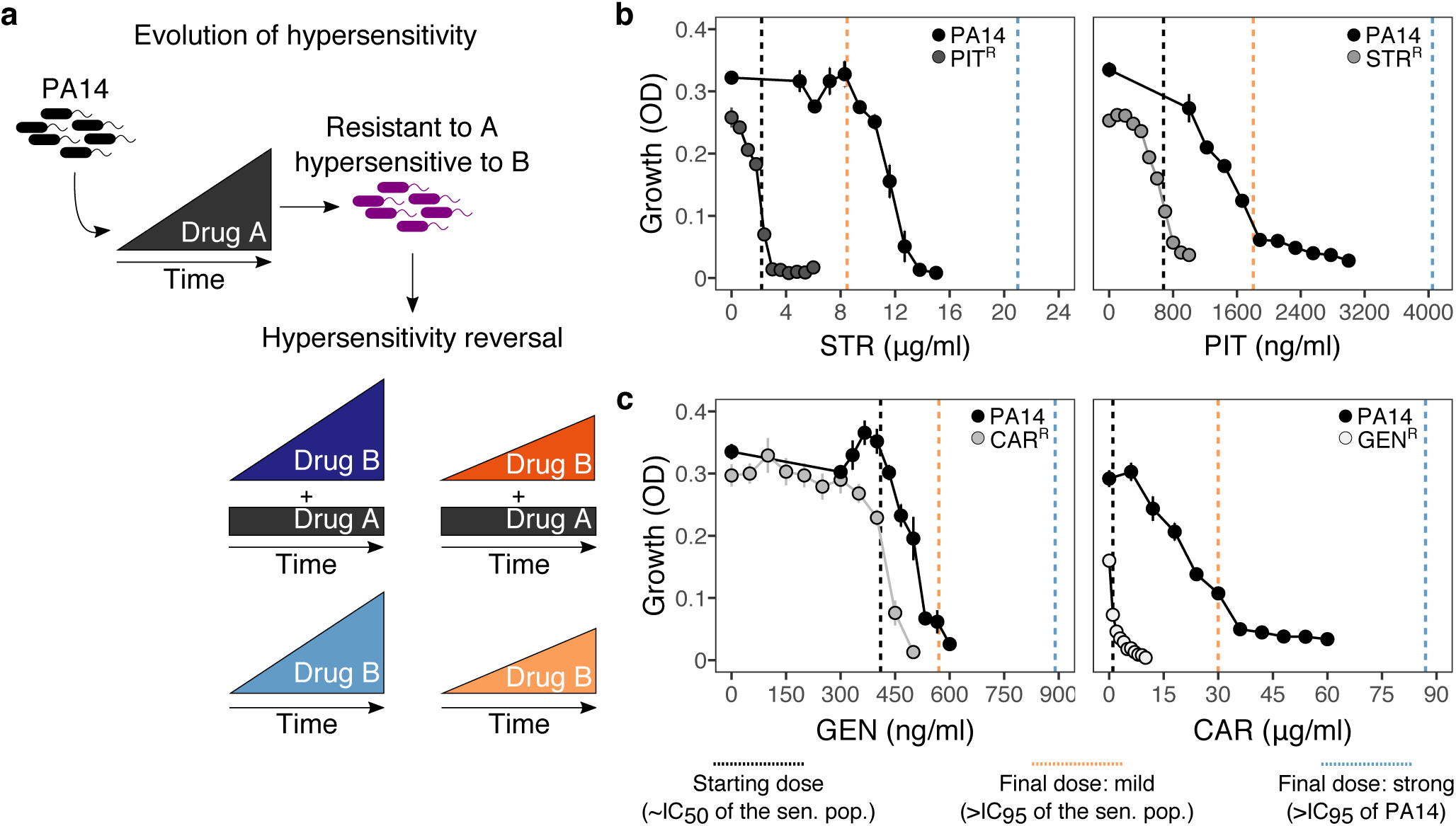
Reciprocal collateral sensitivity and experimental design. **a**, Two-step experimental evolution: resistant populations of *P. aeruginosa* PA14 were previously experimentally evolved (Barbosa et al., 2017) with increasing concentrations of a particular drug (here labelled A), resulting in bacteria becoming hypersensitive to other drugs (here labelled B). In a second step, resistant clones were experimentally evolved in the presence of drug B, using four selection regimes: (i) strong dose increase of drug B in the presence of a constant high dose of drug A; (ii) mild dose increase of B in the presence of A; (iii) strong dose increases of B in the absence of A; and (iv) mild dose increase of B in the absence of A. Concentrations of B were increased using linear ramps starting at IC_50_ (dashed black lines) and ending at levels above the IC_95_ of the collaterally sensitive clone (mild increases, dashed orange line), or that of the PA14 wildtype strain (strong increases, dashed black lines; detailed information on concentrations in Supplementary Table 1). **b**, Validated reciprocity of collateral sensitivity for the isolated resistant clones and drug pair PIT/STR, and **c**, CAR/GEN. Mean ± CI95, n=8. CAR, carbenicillin; GEN, gentamicin; STR, streptomycin; PIT, piperacillin with tazobactam; superscript R denotes resistance against the particular drug. Vertical dashed lines indicate the starting (black) and final doses of the mild (light orange) and strong drug increases (light blue).

### Extinction rates were high even under mild selection regimes

The imposed antibiotic selection frequently caused population extinction (Fig. 2 – Source Data 2), even though sublethal drug concentrations were used. Extinction events occurred significantly more often when selection for the original resistance was maintained by the presence of both drugs (extinction in constrained treatments *vs*. only one drug, *χ*^*2*^=12.9, *df*=1, *P*<0.0001; Fig. 2). In treatments with only the second drug B, extinction occurred significantly more often under strong, but not mild concentration increases (strong vs. mild increases in unconstrained environments, *χ*^*2*^=5.5, *df*=1, *P*=0.019). Drug switches with the antibiotic pair STR/PIT was particularly successful, with 33 extinction events (~51%, Fig. 2). The results differed for the CAR/GEN pair, which produced only 7 extinctions (~10%), all restricted to one drug order, GEN->CAR, suggesting asymmetry in the ability to counter collateral sensitivity. From this, we conclude that strong genetic constraints against an evolutionary response to collateral sensitivity caused frequent population extinctions for STR/PIT switches, whereas evolutionary rescue was possible for the GEN/CAR pair, although influenced by drug identity and order.

**Figure 2.**
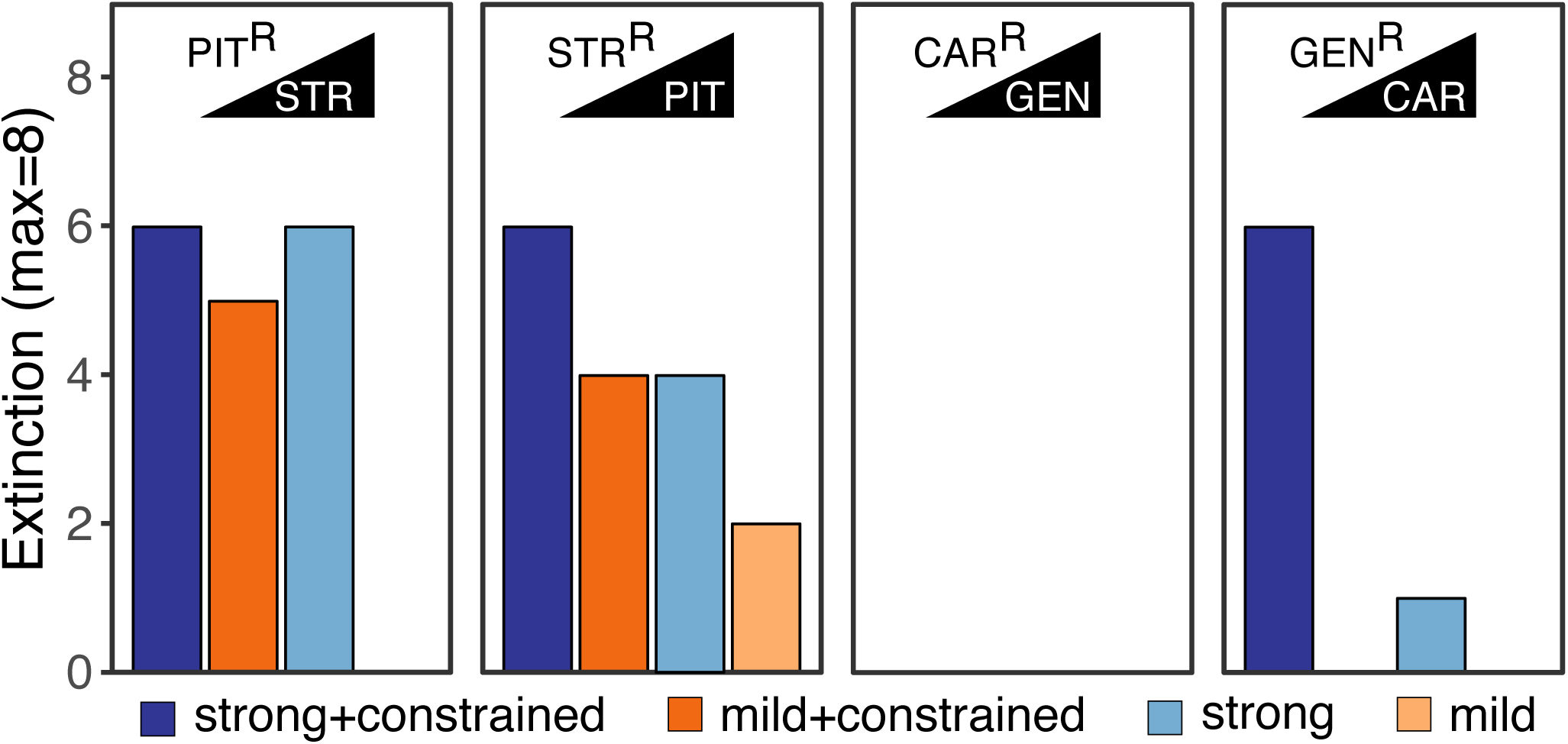
Extinction events during second step of experimental evolution. From left to right, extinction events for PIT^R^-populations adapting to STR, STR^R^-populations adapting to PIT, CAR^R^-populations adapting to GEN, and GEN^R^-populations adapting to CAR. Superscript R indicates resistance against the particular antibiotic, as evolved during the first step of the evolution experiment.

### Novel resistance evolved rapidly in many of the surviving populations

We subsequently focused our analysis of the evolutionary dynamics on the surviving populations of the CAR/GEN pair and identified rapid adaptive responses, especially when not constrained by the presence of the two drugs (Fig. 3). This analysis was not possible for the STR/PIT pair, because few populations survived treatment. For the CAR/GEN pair, we measured bacterial adaptation using relative biomass (see methods and Roemhild et al., 2018) and found it to have increased in all surviving populations (Fig. 3a and b – Source Data 3). For both drug orders, the increase was significantly slower in the constrained treatments, and, to a lesser extent, for the strong concentration increases (Fig. 3a and b, Supplementary Table 2). In consistency with the asymmetry in extinction, the CAR->GEN switch (with no extinction) maintained a high relative biomass across time, while the reverse direction GEN->CAR (with extinction) produced lower relative biomass levels. These results indicate that *P. aeruginosa* can evolve resistance against a drug, to which it had previously shown hypersensitivity, and that such evolutionary rescue is favored for the suboptimal switch. We next asked how the new adaptation influenced the original drug resistances.

**Figure 3.**
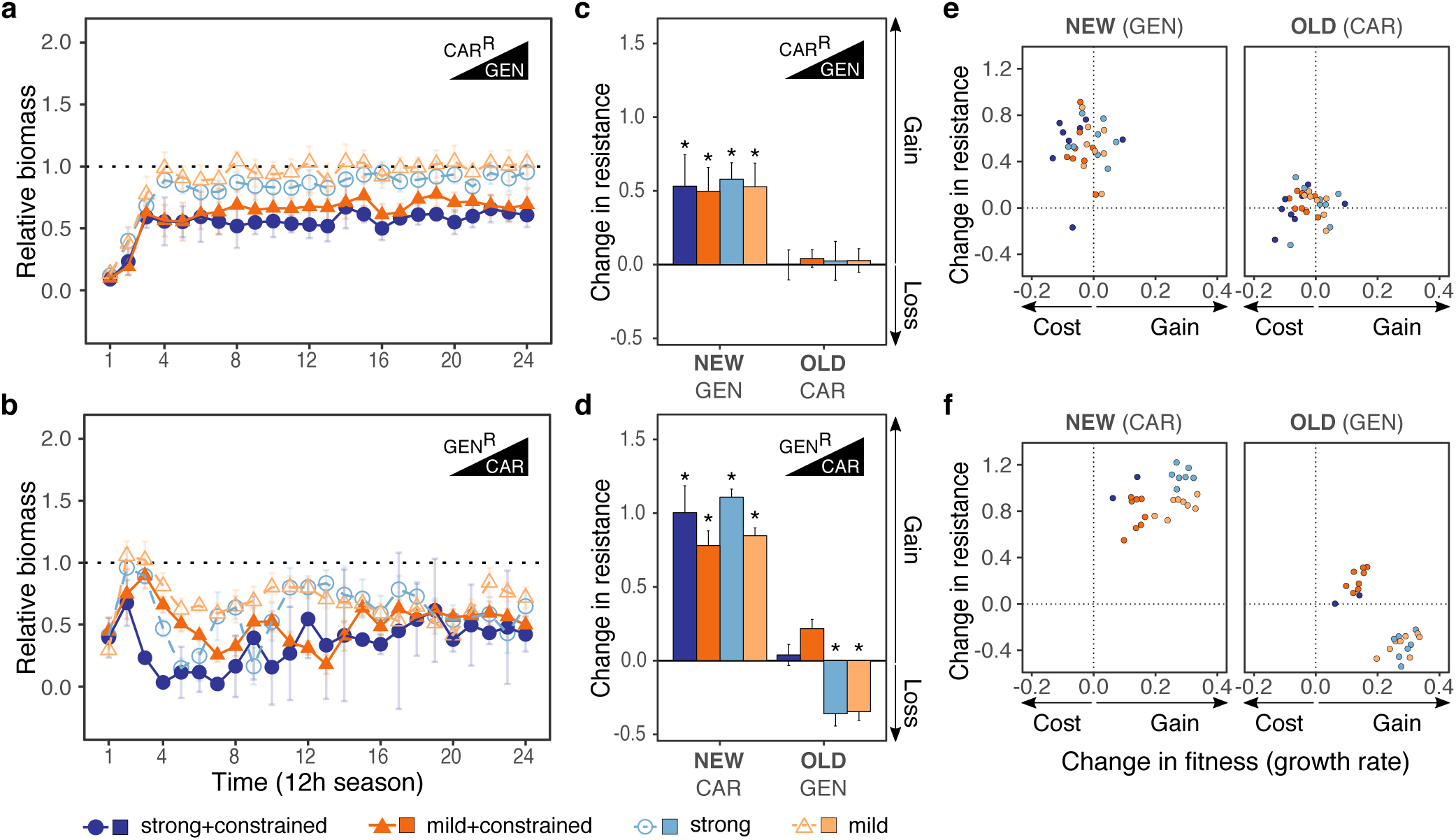
Contrasting evolutionary stability of collateral sensitivity for CAR->GEN and GEN->CAR switches. Evolutionary dynamics of surviving populations expressed as relative biomass for **a**, CAR^R^-populations during selection with GEN, and **b**, GEN^R^-populations during selection with CAR. The dotted horizontal line indicates growth equal to untreated controls. Mean ± CI95, number of biological replicates differs due to extinction (min=2, max=8). Changes in antibiotic resistance at the end of the second-step evolution experiment for **c**, CAR^R^-populations after selection with GEN and **d**, GEN^R^-populations after selection with CAR. Resistance was tested either against the drug towards which bacteria initially expressed resistance after the first evolution experiment (indicated as OLD), or the drug used during the second experiment (indicated as NEW). The change is measured by cumulative differences in dose-response before and after the second evolution experiment (i.e., the original antibiotic resistant clone *versus* its evolved descendants). Mean ± CI95, n = 2-8 biological replicates (differences due to extinction). Asterisks indicate significant changes in resistance (one-sample *t*-test, µ=0, FDR-adjusted probabilities). Correlation between fitness and resistance changes for **e**, CAR^R-^->GEN-lineages and, **f**, GEN^R^->CAR-lineages. The change in fitness (X-axis) is inferred from the growth rate under no-drug conditions of the evolved population at the end of the second evolution experiment relative to that of the respective starting clone, derived from the first evolution experiment (obtained from Barbosa et al., 2017). The corresponding changes in resistance are as shown in c and d. CAR, carbenicillin; GEN, gentamicin; superscript R denotes resistance. Superscript R denotes resistance against the particular drug.

### Drug order determined re-sensitization or emergence of multidrug resistance

Adaptation in the surviving populations of the CAR/GEN pair caused multidrug resistance in the suboptimal switch, but re-sensitization to similar levels than the PA14 ancestor (Supplementary Fig. 1 – Source Data 4 and 5) in the alternative switch (Fig. 3c and d – Source Data 6). In detail, all surviving populations significantly increased resistance against the second drug (Fig. 3c and d; Supplementary Table 3) – in agreement with the recorded biomass dynamics. In the suboptimal switch, CAR->GEN, all populations maintained their original resistance, thereby yielding bacteria with multidrug resistance. This was different for the alternative direction GEN->CAR, where the original resistance was only maintained when both drugs were present in combination (constrained environments). Only under unconstrained evolution, we observed cases of significant re-sensitization to the first drug. We conclude that drug order can determine treatment efficacy, enhance or minimize multidrug resistance, and, in specific cases, lead to a re-sensitization towards the first drug in the surviving populations, as required for applicability of collateral sensitivity cycling (Imamovic & Sommer, 2013).

We hypothesized that the contrasting evolutionary outcomes in constrained *versus* unconstrained treatments of the GEN->CAR switch were caused by an additional trade-off, in this case between drug resistance and growth rate. The starting clones for the second evolution experiment had significantly impaired growth rate and final yield under drug-free conditions, with respectively ~25-50% and 10-50% reductions in fitness relative to the ancestor (Barbosa et al., 2017). As a consequence, selection may have favored variants with higher growth rates. For this particular switch, we indeed found increased growth relative to the resistant ancestral clone for all treatments. The growth rate increases were significantly larger in lineages from unconstrained than constrained treatments (Fig. 3f – Source Data 7, Supplementary Fig. 2 – Source Data 7). In this case, resistance levels also increased slightly (but not significantly) for the unconstrained compared to the constrained treatments (Supplementary Table 4). Similar variations were not observed for the alternative switch CAR->GEN (Fig. 3e – Source Data 7, Supplementary Fig. 2, Supplementary Table 4). We conclude that re-sensitization was favored over multidrug resistance in the GEN->CAR unconstrained treatments, because it provided the advantage of increases in growth rate and, to a lesser extent, resistance to the new drug. This data suggests that fitness costs can determine treatment outcome upon collateral sensitivity switches.

### Whole genome sequencing identified possible targets of antibiotic selection

We used population genomic analysis to characterize specific functional changes that were likely targeted by antibiotic selection and allowed populations to survive the second evolution experiment for the CAR/GEN pair (Fig. 4 – Source Data 8). In particular, we sequenced whole genomes of the resistant starting clones from the beginning and 21 surviving populations from the end of the second evolution experiment. The evolution of multidrug resistance in the suboptimal switch CAR->GEN can be explained by the sequential fixation of mutations including, under unconstrained conditions, those in *ptsP* (Fig. 4a – Source Data 6), a main component of the global regulatory system of ABC transporters and other virulence factors (Feinbaum et al., 2012). Similarly, under constrained conditions, we found mutations in the NADH-dehydrogenase genes *nuoD or nuoG* (Fig. 4a), which are known to influence proton motive force and resistance against aminoglycosides upon inactivation (El’Garch, Jeannot, Hocquet, Llanes-Barakat, & Plésiat, 2007).

**Figure 4.**
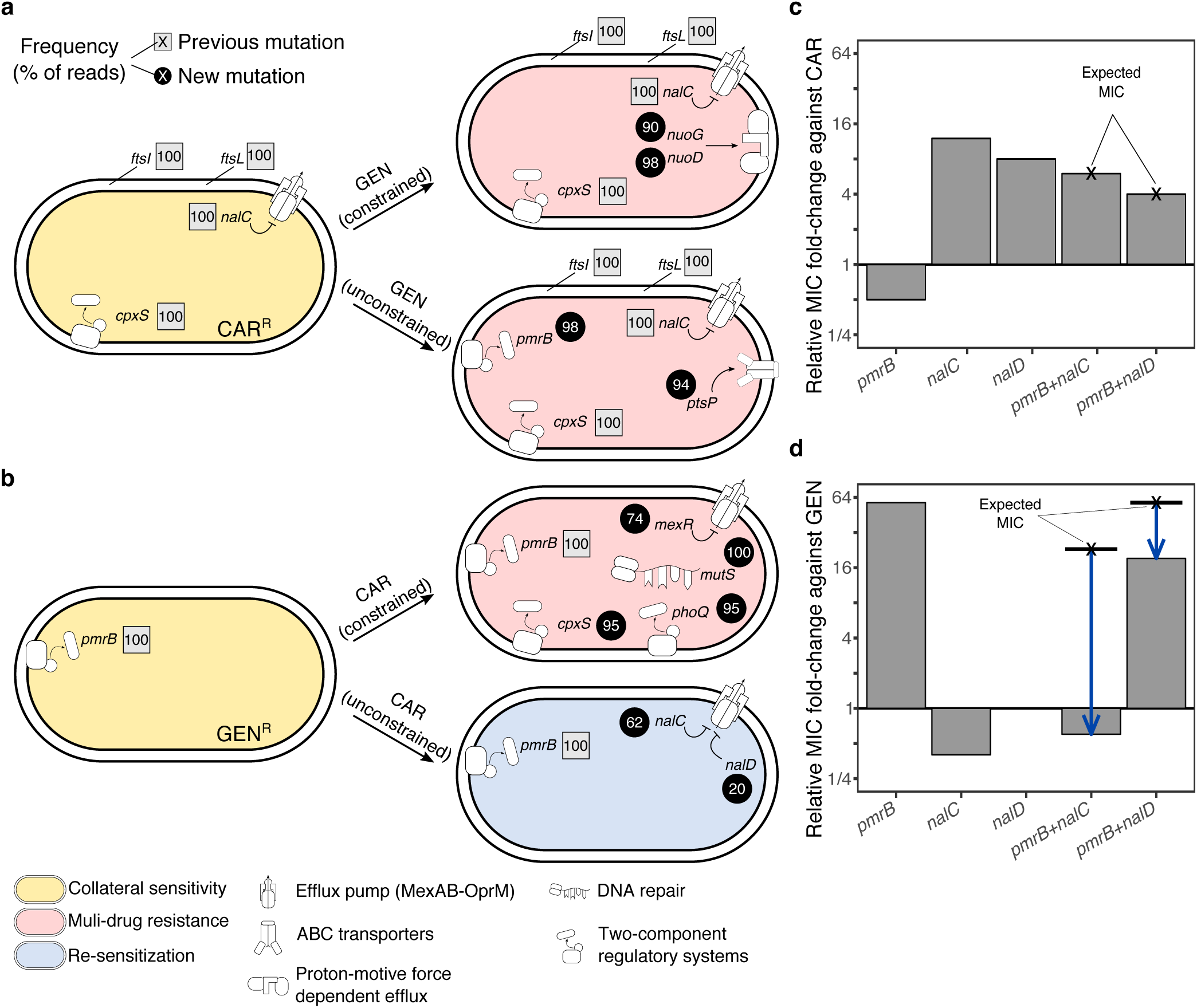
Genomics of experimental evolution for the CAR/GEN drug pair. **a**, Most relevant genomic changes in CAR-resistant populations selected with GEN, and **b**, GEN-resistant populations selected with CAR. Square symbols next to gene names indicate ancestral resistance mutations (obtained from Barbosa et al., 2017), and circles indicate newly acquired mutations. The numbers inside these symbols indicate variant frequencies (as inferred by the percentage of reads in population genomics data) and correspond to the lowest frequency found among the sequenced populations from the respective treatment. The evolved resistance phenotype is highlighted by color shading (see legend in bottom left). All mutations are listed in Source Data 8. **c**, MIC relative to PA14 against CAR and **d**, GEN, of single and double mutant strains. The cross and bold horizontal line indicates the MIC as expected by the individual effects. Blue arrows highlight epistatic effects.

For the more effective switch GEN->CAR, multidrug resistance in the constrained treatments coincided with mutations in *mexR, phoQ, and cpxS*, an independent regulator of MexAB-OprM (X.-Z. Li, Elkins, & Zgurskaya, 2016) and two-component regulators involved in aminoglycoside resistance (Gooderham & Hancock, 2009) and envelope stress response (Roemhild et al., 2018), respectively. The re-sensitization towards the first drug in the unconstrained GEN->CAR treatments was associated with two main types of mutational changes at high frequencies across several replicates, including (i) mutations in *nalC* and *nalD* that upregulate the expression of the multidrug-efflux system MexAB-OprM in *P. aeruginosa* (X.-Z. Li et al., 2016); and (ii) large deletions in *pmrB*, which is part of a two-component regulatory system (Fig. 4b – Source Data 6). Mutations in *nalC* were previously shown to mediate both resistance to CAR and hypersensitivity to GEN (Barbosa et al., 2017). Thus, re-sensitization to GEN may be caused by antagonistic pleiotropy of *nalC* mutations that override the resistance of the original *pmrB* mutation (Source Data 8). In addition, there may be epistasis between the two functional modules. A complementary mechanism for re-sensitization against GEN is the re-mutation of *pmrB* (Source Data 8). In three cases *nalC* mutations coincided with mutations in *pmrB*, including two deletions of 17 and 225 base pairs. Whilst the original SNP in *pmrB* altered gene function, the latter deletions may have suppressed the expression of the original SNP by pseudogenizing the gene. We conclude that mutations in the *nalC or nalD* regulators of the MexAB-OprM pump, sometimes in combination with follow up mutations in *pmrB* are likely to account for the re-sensitization towards the first drug GEN.

### Functional genetic analysis revealed asymmetric epistasis among adaptive mutations

We next investigated whether epistasis between the two functional modules of efflux regulation (MexAB-OprM regulation by *nalC* or *nalD*) and surface charge modification (*pmrB*) may have contributed to re-sensitization using functional genetic analysis. The respective single and double mutations were re-constructed in the common ancestral background of PA14 (see methods for the specific mutations) and changes in resistance against CAR and GEN were measured using fold change of minimal inhibitory concentrations (MIC, Fig. 4c and d – Source Data 9). On CAR, the *pmrB* mutant had half of the MIC of PA14 (confirming collateral sensitivity), whilst *nalC* and *nalD* mutants had increased resistance to CAR. The double mutants had lower MIC on CAR than the *nalC* and *nalD* single mutants (Fig. 4c). The extent of MIC changes in the double mutants corresponded to the product of the individual effects in the respective single mutants, thus indicating an additive interaction among mutations on CAR. On GEN, however, the double mutants had substantially lower MICs than expected from the single mutants (Fig. 4d), strongly suggesting negative epistasis. In detail, GEN-resistance relative to PA14 was 0.4x for *nalC* (collateral sensitivity), 1x for *nalD*, and 57x for *pmrB* (Fig. 4d). The *pmrB, nalD* double mutant had 3x lower MIC to GEN than expected from the individual effects. The *pmrB, nalC* double mutant had >30x lower MIC to GEN than expected from the individual effects, resulting in greater sensitivity than PA14 (Fig. 4d). Altogether, we conclude that re-sensitization to GEN is mediated by antagonistic pleiotropy and negative epistasis.

## Discussion

Collateral sensitivity is a pervasive feature of resistance evolution, but its potential for medical application is currently debated (Nichol et al., 2019; Podnecky et al., 2018; Roemhild & Schulenburg, 2019). Its promise as a treatment focus is that the exploited trade-off is evolutionarily stable and cannot be easily overcome. As a consequence, it should either drive bacterial populations to extinction or minimize the emergence of multi-drug resistance by re-sensitization to one of the antibiotics. We here tested the validity of these key assumptions with the help of evolution experiments and the model pathogen *P. aeruginosa*. We found that the effective exploitation of evolved collateral sensitivity in sequential therapy is contingent on drug order and combination, fitness costs, and also epistatic genetic interactions (Supplementary Fig. 3).

Evolved reciprocal collateral sensitivity generally limited bacterial adaptation. The effect was strongest when the first antibiotic was maintained and a second was added, as reflected by the elevated extinction rates in the constrained environments. This finding may point to a promising, yet currently unexplored treatment strategy, namely single-drug therapy followed by combination therapy, that can maximize exploitation of the evolutionary trade-off. Yet, extinction rates were even high under unconstrained conditions, when drugs were replaced, and in spite of a relatively mild selection intensity, for example an increase in drug concentration from IC_50_ to IC_95_ over a course of 12 days. We observed higher extinction and slower growth improvements in strong, compared to mild drug increases. This finding is generally consistent with previous studies, performed in different context, in which narrowed mutation space upon fast environmental deterioration increased extinction frequencies (Bell & Gonzalez, 2011; Lindsey, Gallie, Taylor, & Kerr, 2013). Interestingly, extinction rates are often not reported as an evolutionary outcome in related studies, possibly because of a different main focus of the study (Yen & Papin, 2017), or because extinction could not be recorded due to the particular experimental set-up (i.e., usage of a morbidostat; (Yoshida et al., 2017)). Considering that antimicrobial therapy usually aims at elimination of bacterial pathogens and extinction frequencies are known from previous evolution experiments to vary among treatment types (Barbosa et al., 2018; Hansen, Woods, & Read, 2017; Roemhild et al., 2018; Torella, Chait, & Kishony, 2010), their consideration should help us to refine our understanding of treatment efficacy.

In the treatments that replaced drugs, evolutionary stability of the resistance trade-off was determined by drug order. Our results for the CAR/GEN pair suggest that this asymmetric stability was determined by variation in either the extent of hypersensitivity and/or the associated physiological fitness costs. The extent of hypersensitivity of the GEN-resistant strain towards CAR was substantially larger than that of the reverse case (Fig. 1c). A similar difference was observed for fitness costs under antibiotic free conditions (see bacterial yield in the no-drug environment on the far left in Fig. 1b and c). It is indeed the GEN->CAR switch (rather than the reciprocal CAR->GEN switch) that produced higher extinction levels (Fig. 2), lower adaptation rates (Fig. 3a and b), and a re-sensitization in the surviving populations (Fig. 3c and d). We conclude that, if the degree of hypersensitivity and/or fitness costs is large, it may be more difficult to counter the strong growth restriction in the presence of the second drug within a limited time frame.

Drug re-sensitization in the unconstrained treatment of GEN->CAR was likely dependent on negative epistasis among pleiotropic resistance mutations. In particular, we found that mutations in *pmrB* and the efflux regulators *nalC* and *nalD* interacted negatively with each other and caused a complete re-sensitization of bacteria that were previously resistant against GEN. While re-sensitization reliably occurred for the GEN->CAR treatment, it did not occur in the reverse case. Similar examples of antibiotic re-sensitization were previously reported for *E. coli* and *P. aeruginosa*, but these relied on different mechanisms. For *E. coli*, repeated alternation between two antibiotics led to re-sensitization as a consequence of clonal interference between variants in two genes, *secD* and/or *basB*. The change between drugs prevented fixation of the competing variants, thus maintaining pleiotropic alleles and thereby the allele causing resistance to one drug and hypersensitivity to the other (Yoshida et al., 2017). In the previous example for *P. aeruginosa*, hypersensitivity to a β-lactam depended on an expression imbalance of the MexAB-OprM and the MexEF-OprN efflux systems after exposure to a fluoroquinolone (Maseda et al., 2004; Sobel, Neshat, & Poole, 2005; Yen & Papin, 2017). Interestingly, partial re-sensitization against the aminoglycoside tobramycin was dependent on adaptive resistance, a phenomenon mediated by the MexXY-OprM efflux pump, whereby expression, and consequently resistance, is induced by the presence of the drug, but then reverted after its removal (Hocquet et al., 2003; Yen & Papin, 2017). We conclude that our finding of negative epistasis between pleiotropic resistance mutations is a previously unknown mechanism underlying re-sensitization. Whilst positive epistasis can substantially amplify resistance gains (Wistrand-Yuen et al., 2018), negative epistasis can limit evolutionary trajectories (Weinreich, Delaney, Depristo, & Hartl, 2006), thus possibly contributing to efficacy of treatment in our case.

We anticipate that the findings of our study could help to guide the design of sustainable antibiotic therapy that controls the infection, whilst reducing the emergence of multidrug resistance. The refined exploitation of collateral sensitivity represents a promising addition to new evolution-informed treatment strategies, including as alternatives specific combination treatments (Barbosa et al., 2018; Chait, Craney, & Kishony, 2007; Evgrafov et al., 2015; Gonzales et al., 2015; Munck et al., 2014), fast sequential therapy (Nichol et al., 2015; Yoshida et al., 2017), or treatments utilizing negative hysteresis (Roemhild et al., 2018).

## Methods

### Material

All experiments were performed with *P. aeruginosa* UCBPP-PA14 (Rahme et al., 1995) and clones obtained from four antibiotic-resistant populations: *CAR-10, GEN-4, PIT-1* and *STR-2* (Barbosa et al., 2017). The resistant populations were previously selected for high levels of resistance against protein synthesis inhibitors from the aminoglycoside family, gentamicin (GEN; Carl Roth, Germany; Ref. HN09.1) and streptomycin (STR; Sigma-Aldrich, USA; Ref. S6501-5G), or alternatively cell-wall synthesis inhibitors from the β-lactam family, carbenicillin (CAR; Carl Roth, Germany; Ref. 6344.2) and piperacillin/tazobactam (PIT; Sigma-Aldrich, USA; Refs. P8396-1G and T2820-10MG). Resistant clones were isolated by streaking the resistant populations on LB agar plates supplemented with antibiotics and picking single colonies after an overnight growth at 37°C. Antibiotic stocks were prepared according to manufacturer instructions and frozen in aliquots for single use. Evolution experiments and resistance measurements were performed in liquid M9 minimal media supplemented with glucose (2 g/l), citrate (0.5 g/l) and casamino acids (1 g/l).

### Measurements of reciprocal collateral sensitivity

The previously reported collateral sensitivity trade-off (Barbosa et al., 2017) was confirmed for this study, by measuring sensitivity of the resistant populations *CAR-10* to GEN, *GEN-4 to CAR, PIT-1* to STR, and *STR-2* to PIT, in comparison to PA14. Populations were grown to exponential phase, standardized by optical density at 600 nm (OD_600_ = 0.08), and inoculated into 96-well plates (100 µl volumes, 5×10^6^ CFU/ml) containing linear concentrations of antibiotics (10 concentrations, 8 replicates each). Antibiotic concentrations were spatially randomized. Plates were incubated at 37°C for 12 h, after which growth was measured by OD_600_ with a BioTek plate reader.

### Experimental evolution

To test the evolutionary stability of reciprocal collateral sensitivity, we challenged clones from previously evolved resistant populations with increasing concentrations of new antibiotics against which the resistant populations showed hypersensitivity (so called collateral sensitivity): *CAR-10* with GEN, *GEN-4* with CAR, *PIT-1* with STR, and *STR-2* with PIT. Stability was assessed with 12-day evolution experiments using a serial transfer protocol (100 µl batch cultures, 2% serial transfers every 12 h; the starting population size for the different populations was approx. 10^6^ CFU/ml), as previously described (Roemhild et al., 2018). Each population was evaluated with 8 replicate populations (4 clones x 2 technical replicates distributed in two plates: plate A and plate B) for each of 5 treatment groups: (i) untreated controls; linearly increasing concentration of hypersensitive antibiotic to a low level (ii) or high level (iii), without maintaining selection for previous resistance (unconstrained evolution); or linearly increasing concentration of hypersensitive antibiotic to a low level (iv) or high level (v), with simultaneous selection for previous resistance (constrained evolution). Concentration increases were started with defined initial inhibition levels of 50% (IC_50_) and concluded when concentrations were above the IC_95_ of the hypersensitive strain (mild increases) or IC_95_ of the wildtype PA14 strain (strong increases), as specified in Supplementary Table 1. Antibiotic selection was applied in 96-well plates and population growth was monitored throughout treatment by continuous measurements of OD_600_ in 15 min intervals (BioTek Instruments, USA; Ref. EON; 37°C, 180 rpm double-orbital shaking). Extinction frequencies were determined at the end of the experiment by counting cases in which no growth was observed after an additional transfer to antibiotic-free media and 24 h of incubation. Surviving evolved populations were frozen at −80°C in 10% (v/v) DMSO, at the end of the experiment.

### Relative biomass

The continuous measurements of optical density during treatment provided a detailed growth trajectory that accurately describes the dynamics of resistance emergence. Relative biomass was defined as total optical growth relative to untreated control treatments, and was calculated by the ratio of the areas under the time-OD curves of treated compared to untreated controls that are passaged in parallel, as previously described (Roemhild et al., 2018).

### Resistance of evolved populations

Resistance of evolved populations was measured for the respective antibiotic pairs (GEN/CAR or STR/PIT), as described above, but using two-fold concentrations (1/4 to 8x the MIC of the starting clone). The respective starting clones of each evolved population served as controls and were measured in parallel. Resistance changes were quantified by subtracting the area under the dose-response curve of the evolved populations from that of the ancestral clones. Positive values indicate that the evolved lineages are more resistant than their ancestor, values close to zero indicate equivalent resistance levels, and negative values denote a loss of resistance. For the cases of re-sensitization against GEN, we performed the experiments a second time including the PA14 ancestor to serve as an additional control (Supplementary Fig. 1).

### Growth rate analyses

Maximum exponential growth rates of evolved and ancestral populations were calculated from growth curves in drug-free media, using a sliding window approach. For measurements, sample cultures were diluted 50x from early stationary phase into 96-well plates (100 µl total volume) and growth was measured by OD_600_ every 15 min for 12 h. Growth rate were calculated from log-transformed OD data for sliding windows of 1 h, yielding two-peaked curves indicating initial growth on glucose and citrate. The reported values the maximum values of the first, larger peak. The values reported in Fig. 3 are the changes of growth rate in evolved populations relative to their resistant ancestors.

### Genomics

We re-sequenced whole genomes of 5 starting clones (CAR-10 clone 2, GEN-2 clones 1-4), and 21 evolved populations (all descendants of these clones from plate B, including 5 untreated evolved control populations and 16 populations adapted to different treatment conditions) using samples from the end of the evolution experiments. Frozen material was thawed and grown in 10 ml of M9 minimal medium for 16-20 h at 37°C with constant shaking. Genomic DNA was extracted using a modified CTAB buffer protocol (Schulenburg et al., 2001) and sequenced at the Institute for Clinical Microbiology, Kiel University Hospital, using Illumina HiSeq paired-end technology with an insert size of 150 bp and 300x coverage. For the genomic analysis, we followed an established pipeline (Jansen et al., 2015). Briefly, reads were trimmed with Trimmomatic (Bolger, Lohse, & Usadel, 2014), and mapped to the UCBPP-PA14 reference genome (available at http://pseudomonas.com/strain/download) using bwa and samtools (H. Li & Durbin, 2010; H. Li et al., 2009). We used MarkDuplicates in Picardtools to remove duplicated regions for single nucleotide polymorphisms (SNPs) and structural variants (SVs). To call SNPs and small SV we employed both heuristic and frequentist methods, only for variants above a threshold frequency of 0.1 and base quality above 20, using respectively VarScan and SNVer (Wei, Wang, Hu, Lyon, & Hakonarson, 2011). Larger SVs were detected by Pindel and CNVnator (Abyzov, Urban, Snyder, & Gerstein, 2011; Ye, Schulz, Long, Apweiler, & Ning, 2009; Ye et al., 2009). Variants were annotated using snpEFF (Cingolani et al., 2012), DAVID, and the *Pseudomonas* database (http://pseudomonas.com). Variants detected in the untreated evolved populations were removed from all other populations and analyses as these likely reflect adaptation to the lab media and not treatment. The fasta files of all sequenced populations here are available from NCBI under the BioProject number: PRJNA524114

### Genetic manipulation

To understand re-sensitization, we analyzed candidate mutations from the GEN->CAR switch. The *nalD* mutation 1551588G>T (resulting in amino acid change p.T11N, as observed in replicate populations b24_G8, b24_D9, and b24_A9) was introduced into the PA14 genetic background using a scar-free recombination method (Trebosc et al., 2016). The same techniques were previously used to construct the mutants *nalC* (deletion 1391016-1391574) and *pmrB* (5637090T>A, resulting in amino acid change p.V136E) in the PA14 ancestor background (Barbosa et al., 2017). Based on these mutants and with the same techniques, we constructed the double mutants *pmrB, nalD* (pmrB p.V136E + nalD p.T11N), and *pmrB, nalC* (pmrB p.V136E + nalC deletion c.49-249, as observed in population b24_F7). Genetic manipulation and confirmation by sequencing was performed by BioVersys AG (Hochbergerstrasse 60c, CH-4057 Basel, Switzerland).

### Epistasis analysis

Resistance of constructed mutant strains was measured in direct comparison to wildtype PA14, as described above. Relative fold-changes in MIC were calculated from dose-response curves. The expected relative resistance of the double mutants was calculated by multiplication of the mutation’s individual effects, as previously described (Wong, 2017). For example, if mutation A conferred a 2-fold increase in resistance and mutation B conferred a 4-fold increase of resistance, the expected resistance of the double mutant AB would be 2×4 = 8. A deviation from this null model indicates epistasis, which can be either positive (greater resistance than expected) or negative (lesser resistance than expected).

## Acknowledgements

We thank D. I. Andersson, R. Kishony, C. Kost, and V. Lázár for feedback on the manuscript. Genome sequencing was performed by G. Hemmrich-Stanisak and M. Vollstedt from the Institute of Clinical Molecular Biology in Kiel, as supported by the DFG Cluster of Excellence EXC 306 “Inflammation at Interfaces”. This research was funded by the Deutsche Forschungsgemeinschaft (DFG, German Research Foundation) individual grant SCHU 1415/12 (to H.S.) and also under Germany`s Excellence Strategy – EXC 22167-39088401 (Excellence Cluster Precision Medicine in chronic Inflammation; H.S., P.R.), the Leibniz Science Campus Evolutionary Medicine of the Lung (EvoLUNG; H.S., C.B.), the International Max-Planck-Research School for Evolutionary Biology (C.B., R.R.), and the Max-Planck Society (H.S., R.R.).

## Author contributions

C.B., R.R. and H.S. designed research, C.B. and R.R. jointly performed experiments and analyzed the data, C.B., R.R. and P.R. analyzed genomic data. All authors wrote the paper.

## Supplementary File

The Supplementary File contains Supplementary Figures S1-S3 and Supplementary Tables S1-S4

## Source Data Files

### Source Data 1 (separate file)

Source data for Figure 1b and 1c. Mean optical density and CI95 values obtained after 12 hours of growth in minimal media and different antibiotics. The populations tested here include the PA14 wt, and four resistant populations described in Barbosa et al., 2017. Each value is the average of 8 technical replicates per bacterial population.

### Source Data 2 (separate file)

Source data for Figure 2. Count data of extinction events. Extinct populations were determined by their inability to grow in rich media after 24 hours of incubation at 37°C.

### Source Data 3 (separate file)

Source data for Figure 3a and 3b. Evolutionary dynamics summarized by the area under the curve (AUC) relative to the treatment with no drugs for the evolution experiments CAR^R^->GEN and GEN^R^->CAR.

### Source Data 4 (separate file)

Source data for Supplementary Figure S1a. Growth characteristics measured by optical density under various drug concentrations for the populations adapted to unconstrained environments (strong and mild), as well as the PA14 wt against gentamicin. Each population-drug concentration was evaluated in triplicate.

### Source Data 5 (separate file)

Source data for Supplementary Figure S1b. Change in resistance of populations adapted to unconstrained environments (strong and mild) relative to the PA14 wt against gentamicin. This was inferred by calculating the difference between the evolved populations and the PA14 wt in the area under the curve across drug concentrations.

### Source Data 6 (separate file)

Source Data for Figure 3c, 3d, 3e, and 3f. Dose-response curves data of surviving populations and the respective ancestors challenged with carbenicillin or gentamicin. Optical density values were recorded after 12 hours of incubation at 37°C.

### Source Data 7 (separate file)

Source Data for Figure 3e, 3f, and Supplementary Figure S2. Growth rate estimates of the surviving populations and the respective ancestors challenged with carbenicillin or gentamicin. Growth rate was calculated as indicated in the Methods section.

### Data Source 8 (separate file)

Source Data for Figure 4a and 4b. Genetic changes compared to *Pseudomonas aeruginosa* PA14 wt strain as determined by whole-genome resequencing (Illumina MiSeq2×150bp PE, Nextera libraries). Isolates are coded with AA-BB-CC-: AA, antibiotic to which they are originally resistant; BB, antibiotic to which the clone shows collateral sensitivity; CC, well in the plate during experimental evolution.

### Data Source 9 (separate file)

Source Data for Figure 4c and 4d. Estimated MIC values for several constructed mutants against carbenicillin and gentamicin.

## Supplementary Information for

### This PDF file includes

Supplementary Figures 1 to 3

Supplementary Tables 1 to 4

Captions for source data files 1 to 9

### Other supplementary materials provided as separate files

Source data files 1 to 9

## Supplementary Figures

**Supplementary Figure 1.**
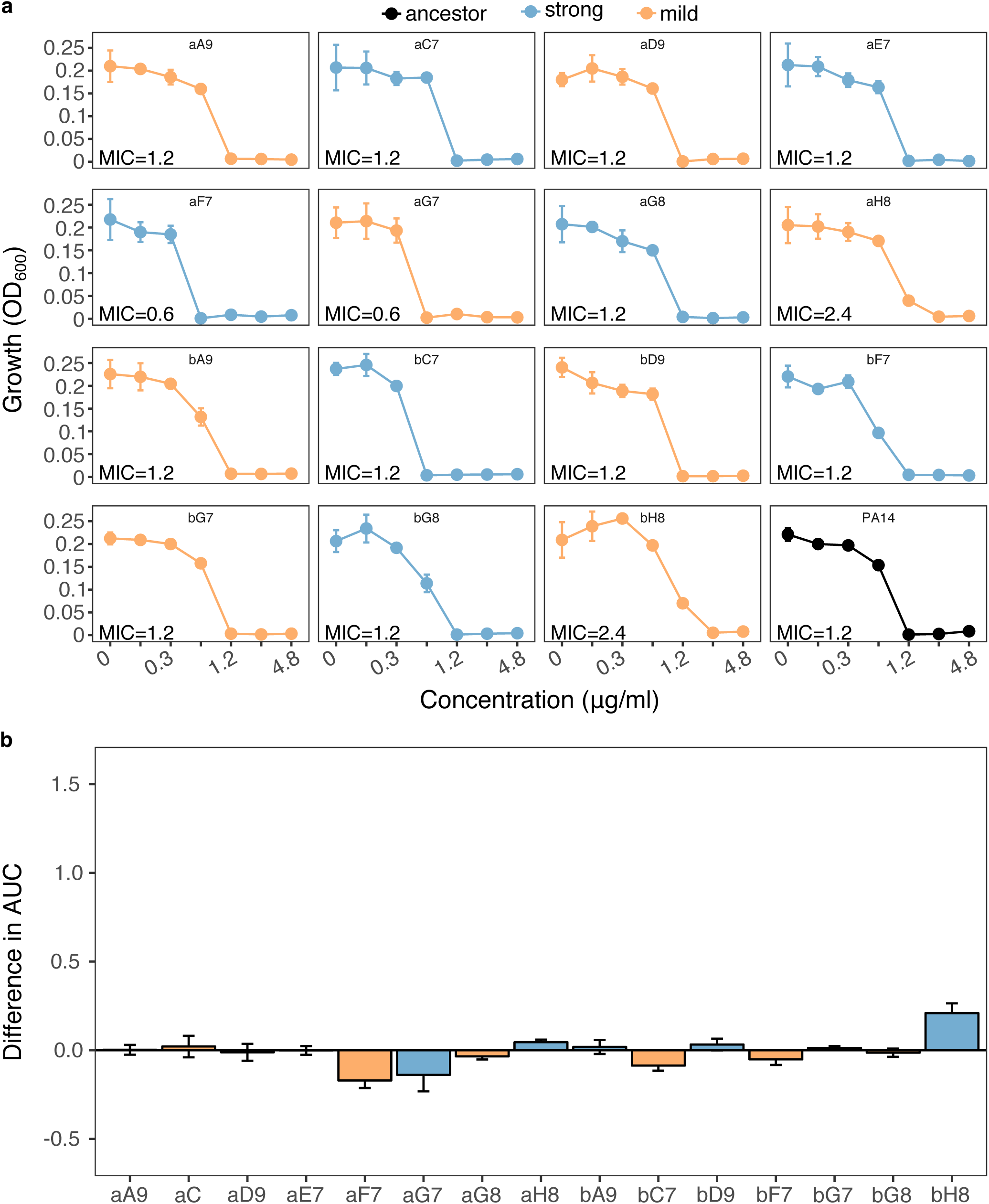
Re-sensitization to gentamicin (GEN) upon adaptation to carbenicillin (CAR). We calculated **a**, dose-response relationships against GEN of 15 populations adapted to strong (n=7, light blue) and mild (n=8, light orange) drug increases compared to the PA14 ancestor (black, bottom-right panel). Mean ± CI95, n = 3 technical replicates. In most cases, the evolved population had the same MIC as PA14. Two populations (aF7 and aG7) showed lower MICs than PA14, while two (aH8 and bH8) showed slightly higher ones. The labels within each graph correspond to the code used during experimental evolution. Data from Source Data 4 **b**, Difference in the area under the curve (AUC) between each evaluated population and the PA14 ancestor. Scaling of the y-axis is equivalent to Fig. 3d. None of the populations was significantly different from the ancestor (Wilcoxon’s test, n=3, adjusted *P* values min > 0.4, and max < 0.9). Data from Source Data 5

**Supplementary Figure 2.**
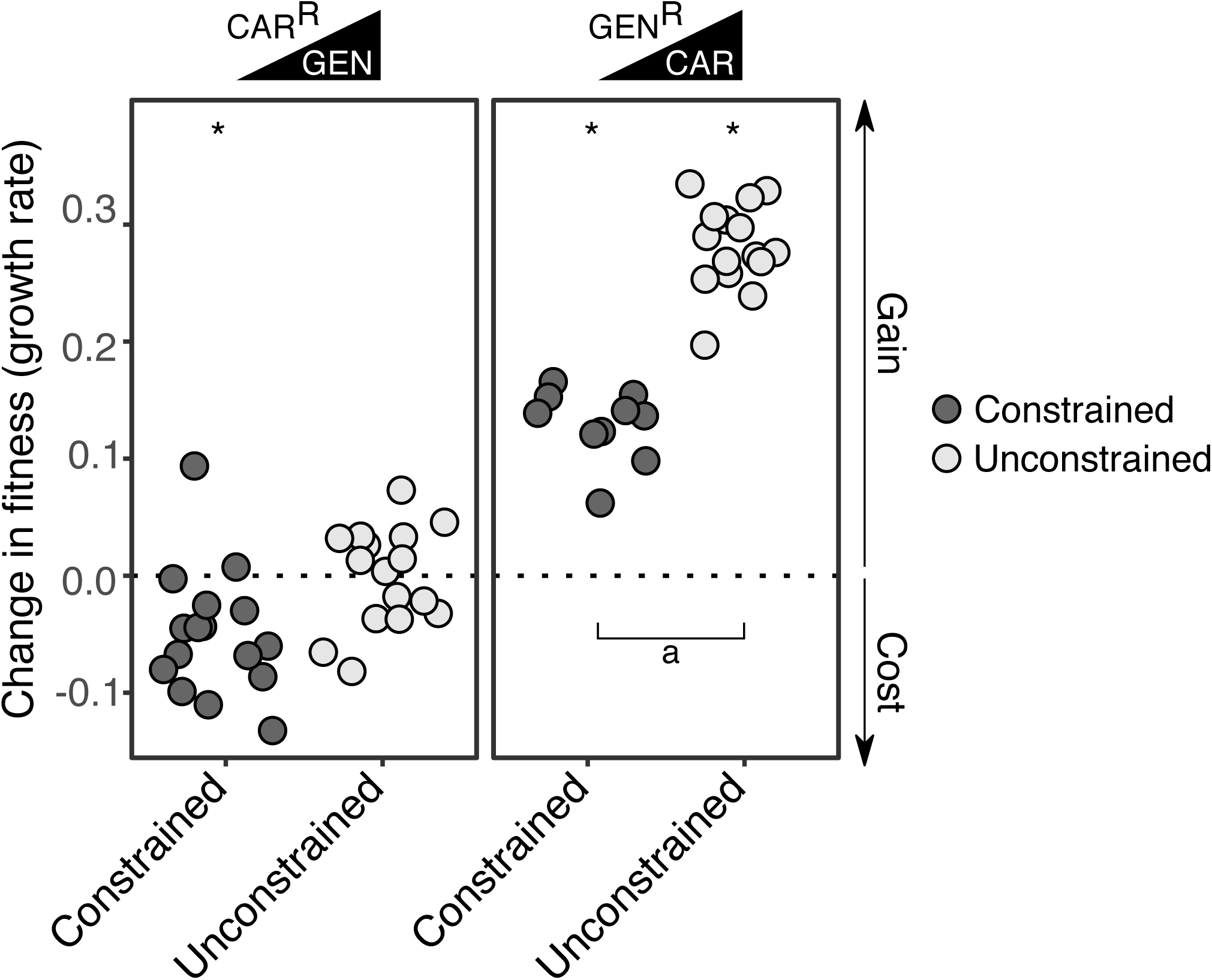
Changes in fitness after experimental evolution. We calculated changes in fitness of the evolved CAR^R^ (left panel) and GEN^R^-populations (right panel) relative to the maximum growth rate of the starting resistant populations (obtained from ref. 8 in the main text). Populations were grouped by whether adaptation was constrained (dark grey) or not (light grey) by the presence of the drug the populations were originally resistant to. Asterisks indicate significant increases or decreases in fitness (One-sample t-test, µ=0, *P* < 0.002). Number of populations per group and experiment vary due to extinction (min=10, max=16). We found significant differences (indicated by the letter a) between constrained and unconstrained treatments in both directions (Two-sample t-test, *P* < 0.0085). Data from Source Data 7.

**Supplementary Figure 3.**
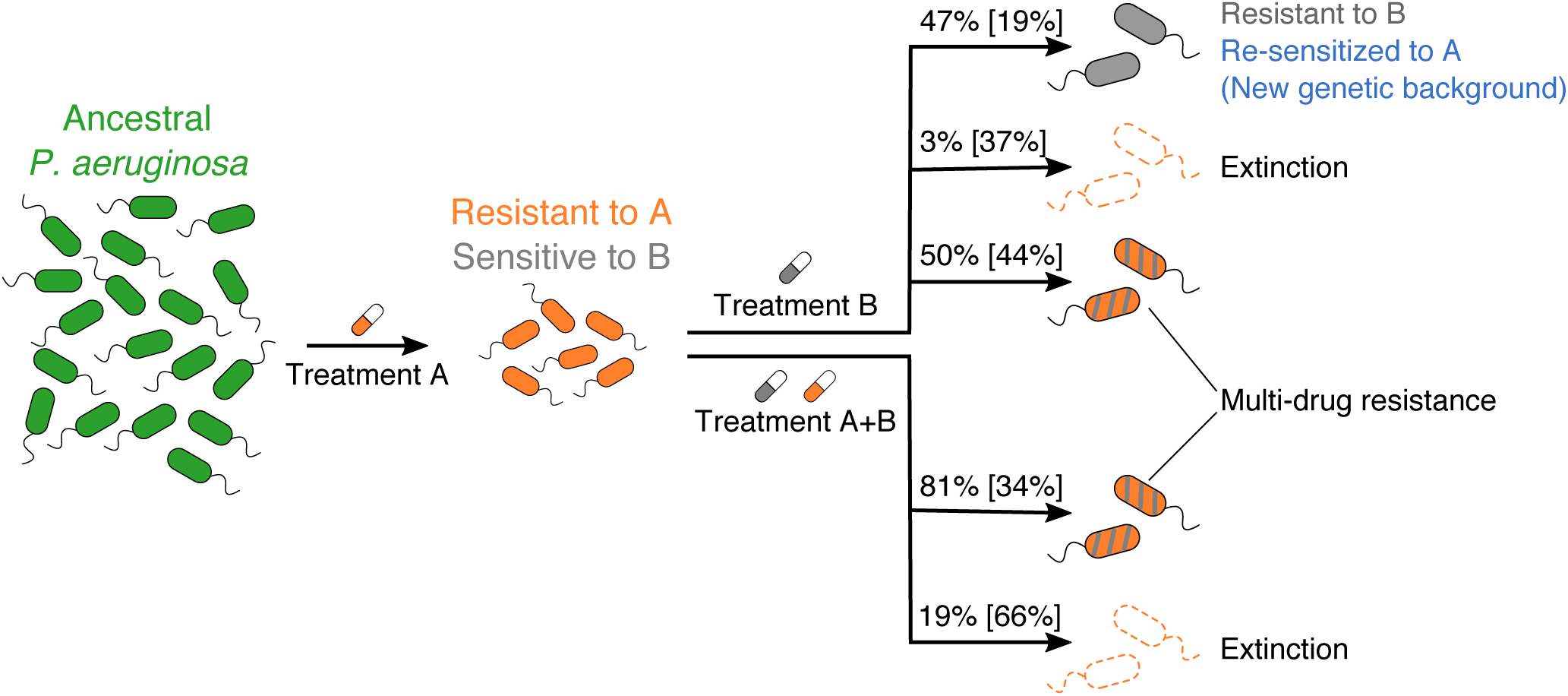
Collateral sensitivity and its evolutionary stability upon antibiotic switches. As *P. aeruginosa* evolves collateral sensitivity after adaptation to one drug (here labeled Treatment A), the subsequent use of other antibiotics (Treatment B, or Treatment A+B) can have several evolutionary outcomes. As now shown by us, the hypersensitive population can evolve resistance to the second drug without modifying the mechanism conferring resistance to the first one, thereby causing multidrug resistance. If the mutations required to escape the sensitivity trade-off are incompatible with the present resistance mechanism, exposure to the second drug could lead to extinction. Alternatively, pleiotropic effects can lead to a situation in which *P. aeruginosa* becomes re-sensitized to the first drug but evolves resistance to the second one. If the second drug is added to the first drug (constrained treatments A+B), then this increases the likelihood of eradicating the bacterial population, but may also come at the risk of multidrug resistance evolution. The percentage of cases from our experiments resulting in each of the described scenarios is shown on top of each arrow. The percentages in black indicate the outcomes for the CAR/GEN experiments, and in cyan those observed for the PIT/STR experiments. A total of 32 replicates is accounted for each possible treatment (with B alone, or with A+B).

**Supplementary Table 1.** Antibiotic concentrations for evolution experiment.

| Previously evolved resistant population | New antibiotic |  |  | For maintenance of original resistance<sup>c</sup> |
| --- | --- | --- | --- | --- |
|  | First dose (~IC<sub>50</sub>) | Final dose mild<sup>a</sup> | Final dose strong<sup>b</sup> |  |
| CAR-10 | 410 ng/ml<br>GEN | 570 ng/ml<br>GEN | 890 ng/ml<br>GEN | +87 µg/ml<br>CAR |
| GEN-4 | 1.0 µg/ml<br>CAR | 30 µg/ml<br>CAR | 87 µg/ml<br>CAR | +890 ng/ml<br>GEN |
| PIT-1 | 2.2 µg/ml<br>STR | 8.5 µg/ml<br>STR | 21 µg/ml<br>STR | +4 µg/ml<br>PIT |
| STR-2 | 0.68 µg/ml<br>PIT | 1.8 µg/ml<br>PIT | 4 µg/ml<br>PIT | +21 µg/ml<br>STR |

<sup>a</sup> Approx. >IC<sub>95</sub> of hypersensitive population specified in column 1
<sup>b</sup> Approx. >IC<sub>95</sub> of wildtype PA14
<sup>c</sup> Added to treatment groups mild+constrained, slow+constrained

**Supplementary Table 2.** Evaluation of the effect of the pace of drug increase (mild or strong) and evolutionary constraint (constrained or unconstrained) on cumulative relative growth^**a**^.

| **Antibiotic** | **Variable** | **$\chi^2$** | **Adjusted *P*** |
| --- | --- | --- | --- |
| GEN | Pace | 14.7 | 0.0002 |
|  | Constraint | 158.1 | <0.0001 |
| CAR | Pace | 18.1 | <0.0001 |
|  | Constraint | 53.8 | <0.0001 |

<sup>a</sup> Separate GLMs were performed for each antibiotic used during experimental evolution with the cumulative relative growth of surviving populations as the response variable, and pace of drug concentration increase (strong or mild) and constraint (constrained or unconstrained) as explanatory fixed factors. Starting clonal population was considered as a nested random factor. We used a type-II Wald $\chi^2$ -test to evaluate the effect of these variables. We used the false discovery rate to adjust the *P* values for multiple comparisons.

**Supplementary Table 3.** Evaluation of the changes in resistance^**a**^.

| **Resistant to** | **Challenged with** | **Treatment** | **Number of populations** | **Adjusted *P*** |
| --- | --- | --- | --- | --- |
| CAR | CAR | No drug | 8 | 0.94485 |
| CAR | CAR | Strong | 8 | 0.7736 |
| CAR | CAR | Strong+constrained | 8 | 0.96151 |
| CAR | CAR | Mild | 8 | 0.6425 |
| CAR | CAR | Mild+bound | 8 | 0.29516 |
| CAR | GEN | No drug | 8 | 0.72291 |
| CAR | GEN | Strong | 8 | 0.00013 |
| CAR | GEN | Strong+constrained | 8 | 0.00489 |
| CAR | GEN | Mild | 8 | 0.00124 |
| CAR | GEN | Mild+constrained | 8 | 0.0016 |
| GEN | CAR | No drug | 8 | <0.00001 |
| GEN | CAR | Strong | 7 | <0.00001 |
| GEN | CAR | Strong+constrained | 2 | 0.10237 |
| GEN | CAR | Mild | 8 | <0.00001 |
| GEN | CAR | Mild+constrained | 8 | <0.00001 |
| GEN | GEN | No drug | 8 | 0.03872 |
| GEN | GEN | Strong | 7 | 0.00065 |
| GEN | GEN | Strong+constrained | 2 | 0.61634 |
| GEN | GEN | Mild | 8 | <0.00001 |
| GEN | GEN | Mild+constrained | 8 | 0.00102 |

<sup>a</sup> *P* values were obtained from a series of Student's t-tests per treatment for populations with ancestral resistance against a given antibiotic and evaluated against two drugs. We used the false discovery rate correction method to adjust *P* values for multiple comparisons.

**Supplementary Table 4.** Changes in resistance between constrained and unconstrained adapted populations in the GEN/CAR experiment^**a**^.

| **Resistant to** | **Challenged with** | **Adjusted *P*** |
| --- | --- | --- |
| CAR | CAR | 0.8156 |
|  | GEN | 0.5694 |
| GEN | CAR | 0.0007 |
|  | GEN | <0.0001 |

<sup>a</sup> *P* values were obtained from a series of two-sample Student's t-tests per treatment for populations adapted to constrained *versus* unconstrained environments. We used the false discovery rate correction method to adjust *P* values for multiple comparisons.

